# Mobility-enhanced virus vectors enable meristem genome editing in model and crop plants

**DOI:** 10.1101/2025.11.19.689159

**Authors:** Kuo Tai Chiu, Harrison Higgs, Mauricio Antunes, Yen Tung Lin, Róisín C. McGarry

## Abstract

CRISPR/Cas9 gene editing revolutionized genetics, but its application is often hampered in non-model plants that are recalcitrant or less amenable to standard plant transformation and regeneration methods. Harnessing viruses to convey guide RNAs (gRNAs) directly to the meristem promises to overcome those limitations and accelerate the generation of edited lines in diverse crops. With several RNA viruses, delivery of gRNAs to the meristem is enhanced with the addition of mobile RNA elements. We hypothesized that incorporating distinct RNA secondary structures as candidate mobility factors in the widely used Tobacco rattle virus (TRV) could propel virus delivery for enhanced meristem editing in non-model species. To test this, we engineered TRV vectors to deliver gRNAs targeting visible marker genes, with each virus incorporating unique mobility factors. We determined optimal virus construction for multiplexed meristem editing by first delivering each virus to *Nicotiana benthamiana* plants harboring the *Cas9* transgene. Strikingly different phenotypes were observed among virus treatments, which were confirmed to represent distinct somatic and heritable editing events. We further tested our hypothesis by leveraging these results to edit pennycress (*Thlaspi arvense*), an emerging oilseed crop. Our results demonstrated successful virus delivery of meristem editing to this non-model plant, underscoring the potential of this approach to deliver targeted genome modifications in diverse crops.

**One sentence summary:** Incorporating RNA mobility factors in TRV affects meristem editing in model plants and crops.

## INTRODUCTION

CRISPR/Cas9 gene editing revolutionized genetics, but its application does not readily extend to non-model plants. Many established and emerging crop species are less amenable to transformation and regeneration technologies. Field pennycress (*Thlaspi arvense* L.) is a potential winter annual cover and oilseed crop. Closely related to Arabidopsis and more commercially valuable brassicas (Lee et al., 2024), pennycress has a short life cycle and extreme cold tolerance, traits that work well for double-cropping with corn and soybean plantings. Natural pennycress stands can produce up to 2.2 metric tons of seed per hectare (McGinn et al., 2019). With seeds containing up to 39% oil by dry weight and a high content of unsaturated fatty acids, pennycress is gaining interest across a range of industrial applications, including feed, plastics, biofuels, and lubricants (Moser et al., 2009). However, opportunities to introduce desirable traits and advance pennycress as a crop are limited because it lacks robust tools for genetic manipulation, such as efficient gene editing. The standard *Agrobacterium tumefaciens*-mediated transformation process is slow, and success varies dramatically across different genetic backgrounds (McGinn et al., 2019); the prolonged tissue culture phases risk somaclonal variation; few independent lines are generated; and the protracted generation time undermines gene functional analysis. Although the floral dip transformation is possible in pennycress, its efficiency remains very low, limiting its utility for rapid trait introduction.

Meristem-based gene editing is an exciting advance: by directly editing dividing cells within meristematic regions, all cell lineages derived from the edited cell will harbor the intended modification to create sectors of edited tissues and organs. Ideally, these sectors will lead to edited flowers with edited pollen and eggs and ultimately edited seeds to dramatically accelerate the generation of an edited line. Plant viruses are increasingly adapted as tools to deliver guide RNAs (gRNAs) for meristem-based editing. The bipartite RNA virus, Tobacco rattle virus (TRV), boasts a broad host range and is widely used in virus-induced gene silencing experiments (Burch-Smith et al., 2004), advancing functional analysis in non-model plants (Andres et al., 2017; Chai et al., 2011; Gao et al., 2011; Hileman et al., 2005; McGarry et al., 2016). To adapt this versatile tool for gene editing, TRV RNA 2 was engineered to deliver gRNAs from the *Pea early browning virus* coat protein promoter (MacFarlane and Popovich, 2000), and, with TRV RNA1, was delivered to *Cas9*-expressing *Nicotiana benthamiana* and Arabidopsis (Ellison et al., 2020; Nagalakshmi et al., 2022b). To facilitate the delivery of gRNAs into germline cells, mobile RNA sequences, many of which assume a characteristic secondary structure, such as tRNAs (Čermák et al., 2017; Nagalakshmi et al., 2022b; Xie et al., 2015) or a truncated non-coding segment from the Arabidopsis *FLOWERING LOCUS T* (*mAtFT*) gene (Ellison et al., 2020), were fused to the 3ʹ end of the gRNAs. The addition of *mAtFT* to the gRNAs dramatically enhanced systemic somatic and heritable editing in *Cas9*-expressing *N. benthamiana*, with the resulting progeny harboring a greater proportion of biallelic mutations in target genes compared with the unmodified TRV controls (Ellison et al., 2020). The incorporation of different mobile RNA sequences in TRV affected heritable editing, with *mAtFT* and *tRNA^Ile^* achieving higher editing frequencies in *Cas9*-expressing *N. benthamiana* and Arabidopsis, respectively (Ellison et al., 2020; Nagalakshmi et al., 2022b). Bookending RNA secondary structure in TRV, with the addition of a hammerhead ribozyme preceding the gRNA and followed with *mAtFT* or *tRNA*, increased editing efficiencies in pepper (*Capsicum annuum* L.) compared to viruses incorporating only one of these RNA structures (Kang et al., 2025). Taken together, these findings suggest that additional RNA secondary structures are tolerated in TRV, and that they have varying effects in enhancing the delivery of gRNAs to the meristem to achieve germline edits.

We questioned whether a broad range of mobile RNA sequences could extend meristem editing from TRV to different plants. MicroRNAs (miRNAs) are small, non-coding RNAs that silence endogenous genes to regulate diverse aspects of plant development and responses to stresses. Transcribed by RNA polymerase II, the primary miRNA (pri-miR) transcript folds into hairpin-like structures, which are recognized by endogenous Dicer1-like proteins, and cleaved into mature miRNAs of 21-23 bps (Zhan and Meyers, 2023). Several miRNAs affect systemic responses, suggesting that these small molecules move from cell to cell and are transported long distances through the phloem. One well-characterized example, *miRNA 399* (*miR399*), functions as a long-distance signal regulating phosphate homeostasis (Chiang et al., 2023; Lin et al., 2008; Pant et al., 2008). When phosphate levels are low, *miR399* is expressed in the shoots. This small molecule is transported to the plant root where it suppresses the expression of the *PHO2* repressor, allowing phosphate to be taken up from the soil. As not all *miRNAs* act systemically, intrinsic features, such as structural motifs of the *pri-miR* transcript, are thought to impact long-distance *miRNA* transport.

We questioned whether the RNA secondary structure intrinsic to *pri-miR399* could enhance gRNA delivery from TRV to plant meristems. In this study, we tested candidate mobile RNA sequences to deliver gRNAs to the meristems of model plants and crops. Mobility factors and gRNAs targeting visible marker genes were assembled in a modular fashion for multiplexed editing from TRV, and these viruses were delivered to *Cas9*-expressing *N. benthamiana*. We demonstrated remarkable differences in somatic and heritable editing efficiencies in *N. benthamiana* based on the mobility factors incorporated. We leveraged these findings to extend meristem editing from TRV to pennycress (*Thlaspi arvense*), an emerging oilseed crop. Our results suggest new methods to achieve targeted genome modifications in non-model and crop plants.

## MATERIALS AND METHODS

### Plasmid constructions

A TRV virus-induced gene silencing (VIGS) control was constructed to silence the homeologous genes encoding *magnesium chelatase subunit I* (MgChl) in tetraploid *Nicotiana benthamiana* (McGarry and Ayre, submitted). Briefly, the 500 bps VIGS target sequence was comprised of 249 bps corresponding to bps 572-820 of the coding sequence of Niben101Scf16898g00001.1 only, and 251 bps identical to the coding sequences of both genes, corresponding to bps 926-1263 of Niben101Scf16898g00001.1 and bps 40-290 of Niben101Scf01308g03004.1. The specificity of the NbMgChl VIGS target was confirmed using the SolGen VIGS tool (Fernandez-Pozo et al., 2015). The NbMgChl VIGS sequence was commercially synthesized, incorporating unique XbaI and SacI restriction endonuclease sites at the 5ʹ and 3ʹ ends, respectively (Twist Bioscience, San Francisco, CA, USA). The synthesized fragment was released from the commercial vector by XbaI and SacI restriction digestion and cloned into the same sites of TRV RNA2 vector pYL156 (Burch-Smith et al., 2006), producing pYL156:NbMgChl.

A VIGS control plasmid was designed to silence the pennycress (*Thlaspi arvense*) *MgChl* ortholog. The 300 bps VIGS target sequence, corresponding to bps 2718-3017 of the Thlar.0034s0042.1 coding sequence, was commercially synthesized with unique XbaI and SacI sites incorporated at the 5ʹ- and 3ʹ-ends, respectively (Twist Bioscience, San Francisco, CA, USA). The VIGS sequence was released from the commercial plasmid by XbaI and SacI digestion, and cloned into the same sites of pYL156, yielding pYL156:TaMgChl.

The *tRNA^Gly^* (Xie et al., 2015) and *mAtFT* (Ellison et al., 2020) were synthesized (General Biosystems, Morrisville, NC, USA). The *pri-miR399* sequence, along with *mir-mimic* and *mir-pro* sequences, which introduced rational mutations into *pri-miR399* (**Fig. S1**), were synthesized (Twist Bioscience, San Francisco, CA, USA).

To construct TRV vectors delivering gRNAs, a modified TRV RNA 2 construct harboring the Pea early browning virus coat protein promoter (PEBVp) upstream of a multiple cloning site (pYL156:PEBVp:MCS) was used (**Fig. S2A**, McGarry and Ayre, submitted). The gRNAs, scaffold, and mobility factors were assembled downstream of the PEBVp in a modular fashion, as described in the **Results**. To construct the viruses harboring gRNAs with *tRNA^Gly^* (**Fig. S2B**) or *mAtFT* (**Fig. S2C**) mobility factors, each cassette was PCR-amplified. The three PCR products were pooled, column-purified, and digested with XbaI, SacI, and BbsI for seamless assembly and directional cloning into pYL156-PEBVp-MCS digested with XbaI and SacI. These viruses are referred to as TRV-tRNA^Gly^ and TRV-mAtFT. To construct the viruses harboring gRNAs with pri-miR399, mir-pro, or mir-mimic mobility factors, each cassette was commercially synthesized and incorporated XbaI and BbsI, BbsI, or BbsI and SacI restriction sites (Twist Bioscience, San Francisco, CA, USA) (**Fig. S2D**). Cassettes were released from the commercial plasmids by restriction digestion, assembled, and directionally cloned into pYL156-PEBVp-MCS digested with XbaI and SacI. These viruses are referred to as TRV-miR, TRV-mir-pro, and TRV-mir-mimic.

To construct a plasmid containing the *Cas9* and *DsRed2* fluorescent marker genes for pennycress transformation, we initially assembled the transcription units (TU) consisting of the CaMV 35S (35Sp) promoter:Cas9, Nopaline Synthase (Nosp) promoter:HygR, and 35Sp:DsRed2 using Goldenbraid assembly methods (Sarrion-Perdigones et al., 2011; Sarrion-Perdigones et al., 2013). All TUs contained the Nopaline Synthase terminator sequence and were further combined into the pDGB3α1 plasmid backbone following similar assembly methods, yielding the pSF218 binary plasmid. Individual pUPD2-domesticated genetic components and pDGB3α1 were all sourced from the Goldenbraid 2.0 kit obtained from Addgene (#1000000076).

### Virus inoculations

Plasmids were introduced to *Agrobacterium tumefaciens* GV3101 pMP90 by electroporation. Single colonies were cultured and prepared for inoculation as described (McGarry and Ayre, 2024). *A. tumefaciens* strains harboring TRV RNA 1 and each recombinant TRV RNA 2 were mixed immediately before delivery to plants. For *N. benthamiana* inoculations, each virus was delivered by syringe-infiltration into the abaxial surface of four expanding leaves from each of six replicate plants. For pennycress inoculations, each virus was delivered by syringe-infiltration into the abaxial surface of leaves from seedlings at different ages (i.e., only cotyledons expanded, two true leaves, or six true leaves) or by vacuum infiltration of whole seedlings. Each virus was delivered to, at minimum, six replicate pennycress plants.

### Plant growth conditions

*Nicotiana benthamiana* harboring the *35Sp:Cas9* transgene were generously gifted by Dr. Daniel Voytas (University of Minnesota). Transgenic seeds were densely sown on the surface of coarse-cut potting mix (2 parts Sta-green Fruit, Flower, and Vegetable Potting Soil Mix: 1 part Sta-green Tree and Shrub Garden Soil: and 1 part perlite) in 3.5-inch pots and allowed to germinate in a growth chamber at 25°C with 16h day / 8 h night. Seedlings were transplanted to individual 3.5-inch pots and returned to the same growth chamber. Approximately three weeks later, plants with four to five true leaves were *Agro*-infiltrated with recombinant viruses. Inoculated plants were domed and incubated at room temperature overnight before returning to a growth chamber set at 25°C and 16 h day / 8 h night.

Three fruits were collected from each of the six replicate *N. benthamiana* plants of each treatment. Seeds were surface-sterilized and germinated on 0.5x MS solid medium supplemented with 1% sucrose. Plates were placed in a growth chamber at 25°C with 16h day / 8 h night.

Pennycress cv. Spring 32-10 harboring the *35Sp:Cas9* transgene was generated by *A. tumefaciens*-mediated vacuum transformation using the pSF218 plasmid. Wild type and homozygous T2 plants were used for virus inoculation experiments.

### DNA isolation and preparation of amplicons for sequencing

Leaf samples (< 50 mg) were collected from the green and white sectors on the first and seventh systemic leaves of all *N. benthamiana* plants. DNA was isolated by CTAB (Doyle and Doyle, 1990). The DNA sequences flanking the predicted *NbMgChl* edited sites were PCR-amplified and prepared for amplicon sequencing (Jacobs et al., 2015; Xue and Tsai, 2015). As the expected PCR products were of similar sizes, 266 bps for Niben101Scf16898g00001 and 291 bps for Niben101Scf01308g03004, each was amplified separately. The first PCR reactions used gene-specific primers that incorporated Illumina A5 or A7 index tails (**Table S1**) with Phire Plant Direct PCR Master Mix (Thermo Scientific, Waltham, MA, USA) in 10 µL reactions. PCR products (4 µL) were visualized by agarose gel electrophoresis and standardized based on band intensity. Approximately equivalent amounts of the two *NbMgChl* amplicons were pooled from each biological sample. The second PCR reactions used the pooled DNA as templates with the A500 and A700 series Illumina index primers and Phire Plant Direct PCR Master Mix (Thermo Scientific, Waltham, MA, USA) in 10 µL reactions arranged in a 96-well plate. PCR products (4 µL) were visualized by gel electrophoresis and standardized based on band intensities. Equivalent amounts of all PCR products were pooled and gel-purified (Promega, Madison, WI, USA). The eluted DNA was quantified by nanodrop. Amplicons were sequenced using an Illumina MiSeq 300-cycle nanokit for 1 million reads at the Genomics Core of the University of North Texas Health Science Center. The resulting FASTQ files were processed using the AGEseq algorithm (Jacobs et al., 2015; Xue and Tsai, 2015).

Pennycress leaf samples were collected from green, white, and pale green sectors of leaves, as well as from the leaf at the next node, from TRV:gTaMgChl-mAtFT-treated plants and uninoculated control plants. DNA was isolated as described. Amplicons were prepared with gene-specific primers (**Table S1**) and analyzed as above.

### RNA isolation and expression analyses

Green and white leaf sectors were harvested from the young, unexpanded leaves of three replicate *N. benthamiana* plants per treatment. Samples were immediately plunged into liquid nitrogen and homogenized using a Retsch mill (MM400; Retsch USA, Newtown, PA, USA). RNA was isolated using IBI Isolate (IBI Scientific, Dubuque, IA, USA) as per the manufacturer’s instructions. Total RNA was visualized by gel electrophoresis and quantified by nanodrop. Approximately 500 ng of total RNA was used to generate cDNA with random hexamers, oligo dT23 primers, and Protoscript II (New England Biolabs, Ipswich, MA, USA). The cDNA was diluted ten-fold, and 1 µL was used in 10 µL quantitative PCR reactions using Powertrack SYBR master mix (Thermo Fisher Scientific, Waltham, MA, USA) with a QuantStudio 6 Pro Real-Time PCR System (Thermo Fisher Scientific, Waltham, MA, USA). Reactions were carried out for 40 cycles using a fast cycle program (initial denaturation at 95 °C for 2 min, followed by 40 cycles at 95 °C for 5 s and 60 °C for 30 s) and melt curve analysis (95 °C for 15 s, 60 °C for 1 min and 95 °C for 15 s), as per the manufacturer’s instructions. Gene expression was determined from the green and white tissue sectors from three biological samples, meaning three independent plants, from each virus treatment. Each RT-qPCR reaction had two technical replicates, and technical replicates that deviated by more than one Cq value were removed from further analysis. Two reference genes, *protein phosphatase 2a* and *EF1α*, were quantified in *N. benthamiana* (**Table S1**;(Ellison et al., 2020). Target transcripts were quantified using primers specific to *NbMgChl* and to the 5′ and 3′ regions of TRV RNA 2. Target gene expression was calculated relative to the mean Cq values of both reference genes. The relative expression was calculated by the ΔΔCt method (Livak and Schmittgen, 2001), with *NbMgChl* compared to uninoculated plants and virus expression compared to TRV-treated plants. The variation was expressed as the standard error of the mean.

### Statistical analyses

Statistical analyses were carried out using IBM SPSS Statistics v30.0.0.0 according to the General Linear Model with multivariate ANOVA. Post-hoc tests by Tukey’s HSD (*p* < 0.05) were used to determine significant differences among treatments.

## RESULTS

### Design and assembly of guide RNAs with assorted mobility factors in viral vectors

To identify the *N. benthamiana* orthologs of the chlorophyll biosynthesis gene *magnesium chelatase subunit I* (*MgChl*), we queried the cotton (*Gossypium hirsutum*) magnesium chelatase proteins (Gohir.A06G200700, Gohir.D06G208200, Gohir.A10G030400, and Gohir.D10G031300) against the *Nicotiana benthamiana* proteome (v1.0.1) in BLASTp searches. Two hits and presumed homeologs were identified: Niben101Scf01308g03004.1 (*NbMgChl1*) and Niben101Scf16898g00001.1 (*NbMgChl2*). To test if these could be used as visible markers, we silenced the transcripts using virus-induced gene silencing (VIGS). Approximately 500 bps of sequences specific to both homeologs were introduced as silencing targets in Tobacco rattle virus RNA 2 vector pYL156, producing TRV:NbMgChl. The recombinant virus was delivered via *Agro*-infiltration to wild-type *N. benthamiana* plants, along with TRV and uninoculated controls. As expected, all replicate VIGS plants produced systemically photobleached leaves by ∼ 7 days post-inoculation (dpi) (**Fig. S3C**), and silencing was sustained for the life of the plants. The TRV-treated plants showed damage to the inoculated leaves (**Fig. S3B**), but new growth was green and healthy, as were the uninoculated controls (**Fig. S3A**). This loss-of-function phenotype confirmed that the *NbMgChl* genes would be useful candidates for comparing virus-based gene editing strategies.

We sought to optimize our viral vectors through rigorous testing in *Cas9*-expressing *N. benthamiana*. We designed a series of viral vectors for multiplexed editing of *NbMgChl1* and *NbMgChl2* and tested the effects of different mobility factors in each virus. First, we identified gRNAs within exon sequences of *NbMgChl1* and *NbMgChl2* using Cas-Designer from CRISPR RGen Tools (Bae et al., 2014; Park et al., 2015). Three 20-bp gRNAs, predicted to target the homeologs with high likelihood and having no predicted off-targets, were selected. To ease downstream analyses, the three gRNAs were positioned within < 200 bps in each gene. Second, we incorporated different predicted RNA secondary structures in TRV to compare the effects of putative mobility factors on meristem editing (**Fig. S1**). The impacts of tRNAs (Čermák et al., 2017; Li et al., 2021; Nagalakshmi et al., 2022b; Xie et al., 2015) and the mutated and truncated Arabidopsis *FLOWERING LOCUS T* (*mAtFT*) on meristem editing are well characterized (Ellison et al., 2020). We compared *tRNA^Gly^* and *mAtFT* with the *primary microRNA 399* (*pri-miR399*) sequence. As the prominent secondary structure of *pri-miRNAs* is lost during maturation, we engineered rational mutations to prevent this processing and maintain the secondary structure (*mir-pro*) or to alter its structure to resemble *mAtFT* (*mir-mimic*; **Fig. S3**, **Table S2**). Next, we implemented a modular strategy to simultaneously assemble the same three gRNAs with a unique candidate mobility factor in TRV:PEBVp. Viruses incorporated three cassettes, each consisting of a 20 bp gRNA, scaffold (76 bps), and mobility factor (**Fig. S2**). Our design was limited to three cassettes such that full cargo size, at most 805 bps, was within reasonable limits for TRV size constraints and similar to published reports (Ellison et al., 2020). For the *tRNA^Gly^*- and *mAtFT*-containing viruses, gRNAs were introduced by overlapping PCR with custom oligonucleotides incorporating recognition sequences for seamless assembly and directional cloning in TRV:PEBVp (**Fig. S2A-C**). For the *pri-miR399*, *mir-pro*, and *mir-mimic*-containing viruses, each cassette, incorporating the same restriction endonuclease recognition sites, was commercially synthesized (Twist Bioscience; **Fig. S2D**). Each cassette was released from the commercial vector and directionally cloned into TRV:PEBVp as above. This vector series thus included, in addition to the control TRV:PEBVp and VIGS control TRV:NbMgChl, TRV:PEBVp:gNbMgChl-tRNA^Gly^ (hereafter “TRV:tRNA”), TRV:PEBVp:gNbMgChl-mAtFT (“TRV:mAtFT”), TRV:PEBVp:gNbMgChl-pri-miR399 (“TRV:pri-miR399”), TRV:PEBVp:gNbMgChl-mir-pro (“TRV:mir-pro”), and TRV:PEBVp:gNbMgChl-mir-mimic (“TRV:mir-mimic”). This series of TRV vectors was predicted to yield quantitative insights for optimized delivery of gene editing tools, which could be readily extended to non-model plants.

### Mobility factors differentially impact early somatic editing in the leaves of *35S:Cas9*-expressing *N. benthamiana* plants

The recombinant TRV constructs were prepared for *Agro*-infiltration (McGarry et al., 2016), and delivered to *Cas9*-expressing *N. benthamiana* plants. By 9 dpi, striking and consistent differences were observed in the systemic leaves of TRV-treated plants (**Fig. 1**). All *N. benthamiana* plants treated with TRV:tRNA (*n* = 6) and TRV:mAtFT (*n* = 6) showed extensive photobleached sectors in the systemic leaves, and these phenotypes continued for the life of the plants (**Fig. 1D, E**). In contrast, plants treated with the TRV:pri-miR399 (*n* = 6), TRV:mir-pro (*n* = 6), or TRV:mir-mimic (*n* = 6) viruses showed a few discrete yellow patches on systemic leaves, which occurred less frequently in newer growth (**Fig. 1F-H**). Included as controls, all of the VIGS-(TRV:NbMgChl) treated plants produced uniformly white systemic leaves (*n* = 6). The TRV-treated *N. benthamiana* plants (*n* = 6) showed some necrosis as previously described, and uninoculated plants remained green and healthy (*n* = 6; **Fig. 1A-C**).

**Fig. 1:**
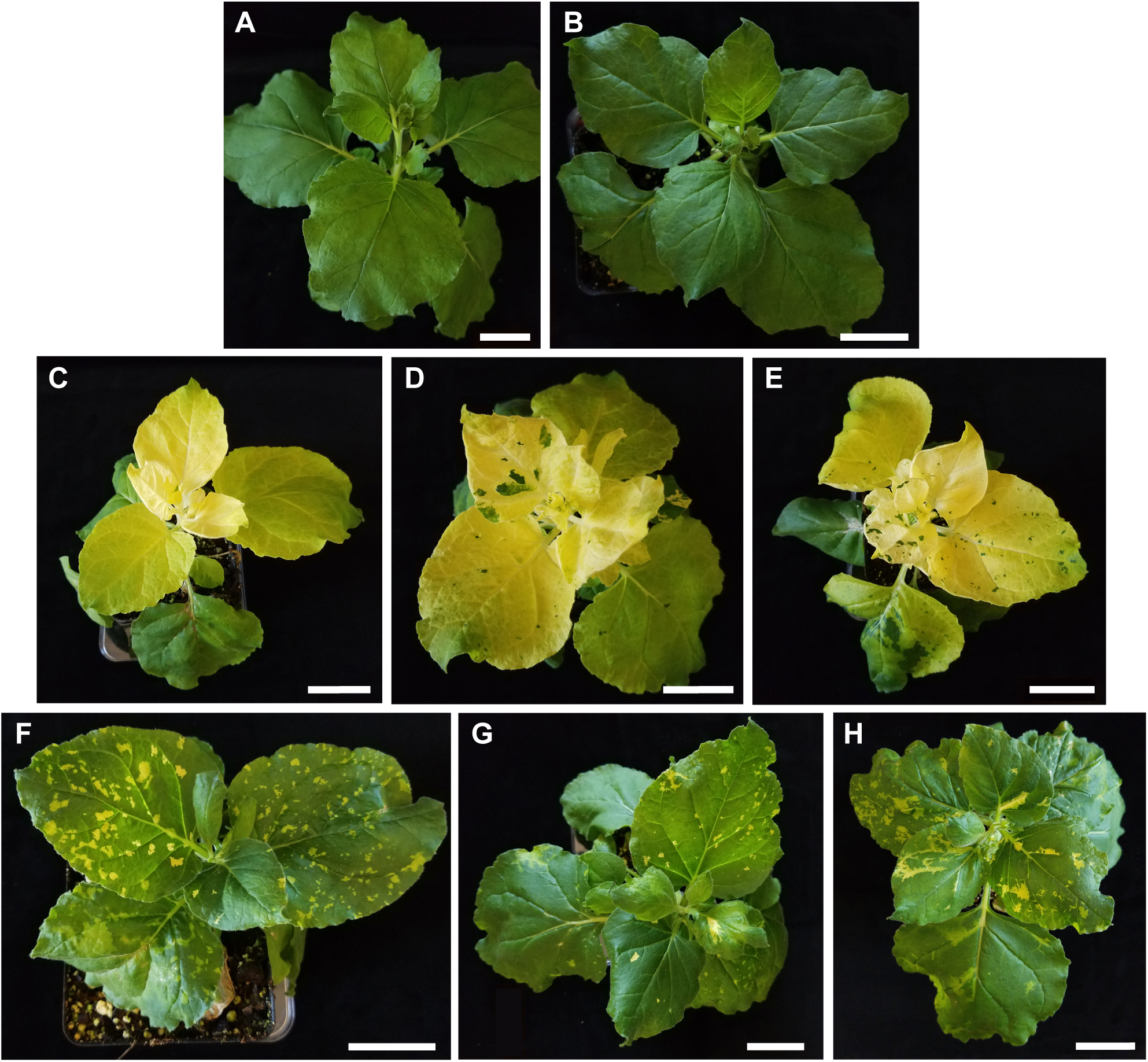
TRV delivers gRNAs targeting the *magnesium chelatase* genes flanked by candidate mobility factors to *Cas9*-expressing *N. benthamiana*. (A) Uninoculated and **(B)** TRV:PEBVp-treated plants remained green. **(C)** Silencing the target genes with TRV:NbMgChl produced systemically photobleached leaves in treated plants. **(D)** TRV:tRNA- and **(E)** TRV:mAtFT-treated plants produced extensive photobleached sectors in systemic leaves, which continued for the life of the plants. **(F)** TRV:miR-, **(G)** TRV:mir-pro-, and **(H)** TRV:mir-mimic-treated plants produced smaller sectors of photobleached tissues, and these sectors occurred less frequently in newer growth. Plants were imaged at 9 dpi. Scale bars are 5 cm.

Because photobleaching was evident in the first systemic leaf, we questioned whether these phenotypes resulted from editing or silencing. To test this, we analyzed the loci for molecular evidence of editing. Genomic DNA was isolated from white and green leaf sectors from the first and seventh systemic leaves from three replicate plants per treatment, and amplicons were prepared for next-gen sequencing. We predicted that editing from any one gRNA could generate small indels, and successful editing at two or more target sites could yield deletions of 18, 95, or 113 bps in each gene (**Fig. 2A**). The loci were considered wild type when 0 - 15% of reads harbored a mutation; monoallelic when the editing efficiency was 35 - 65%; biallelic when 85 - 100% of reads had mutations; and the loci were considered mosaic when 16 - 34% or 66 - 84% of reads contained a mutation (Ellison et al., 2020; Li et al., 2021; Liu et al., 2022).

**Fig. 2:**
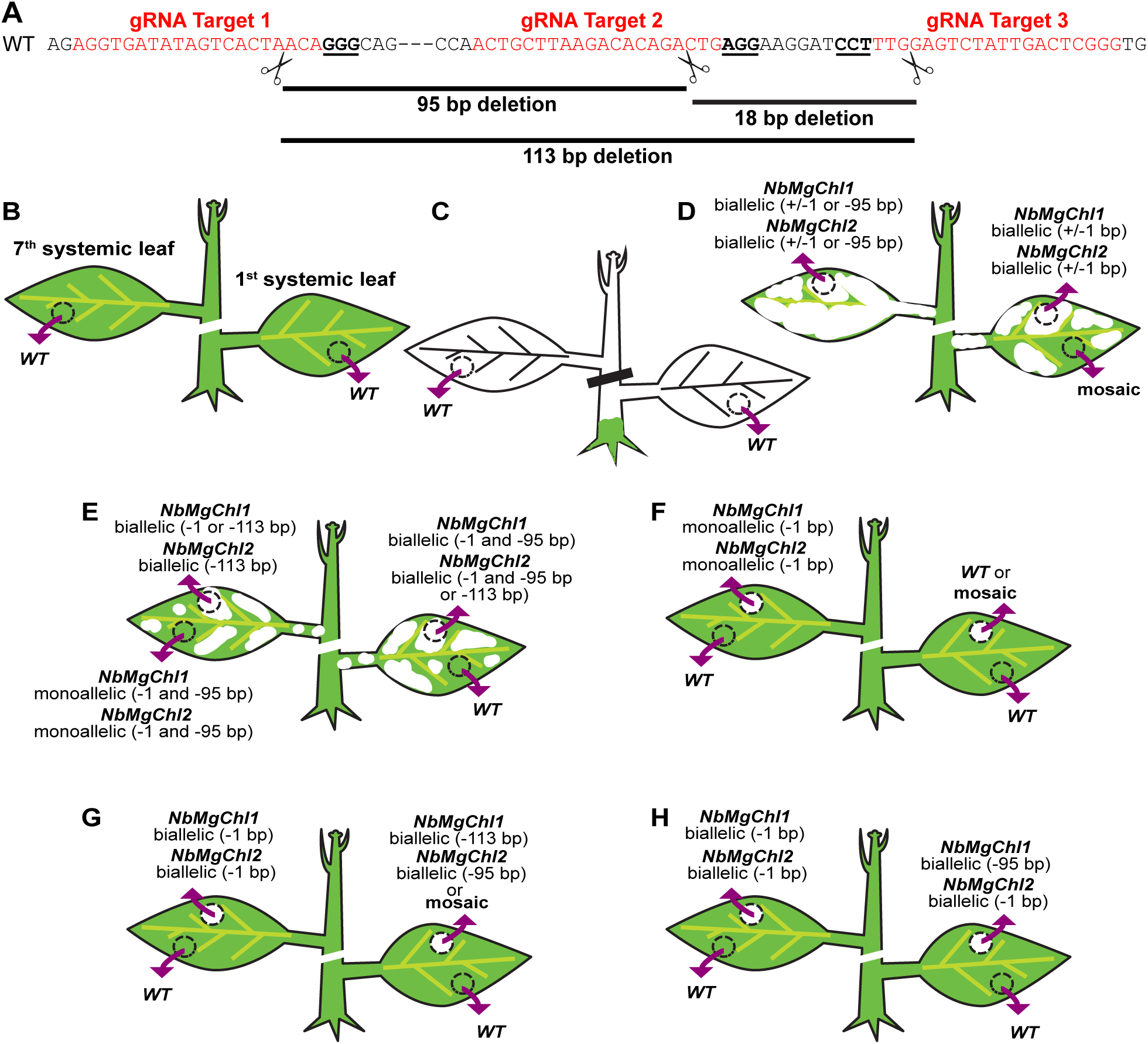
M**o**lecular **evidence of somatic editing. (A)** The *NbMgChl1* and *NbMgChl2* gene sequences are identical in the regions targeted for editing. Three gRNA targets are shown in red font, with the PAM sequence underlined. Scissors represent predicted cleavage sites, with possible deletion sizes from multiple cleavages shown. **(B-H)** Genomic DNA was isolated from the first and seventh systemic leaves and the *NbMgChl1* and *NbMgChl2* loci analyzed by amplicon sequencing. **(B)** The target loci were wild type in the uninoculated and TRV-treated control plants. (C) The target loci were wild type in the systemic white VIGS plants treated with TRV:NbMgChl. TRV:tRNA-treated plants harbored biallelic mutations in both genes in the white tissue sectors. **(E)** TRV:mAtFT-treated plants harbored biallelic deletions in both loci in white tissue sectors and monoallelic deletions in the systemic green sectors. **(F)** TRV:miR-treated plants had monoallelic single-nucleotide deletions in white tissue sectors. **(G)** TRV:mir-pro-treated plants harbored large biallelic deletions early in the infection, but later emerging leaves had smaller single-nucleotide deletions in white tissue sectors. **(H)** TRV:mir-mimic-treated plants harbored larger biallelic mutations early in infection, with later emerging leaves having smaller biallelic mutations in white tissue sectors.

Amplicon sequencing revealed successful and targeted editing of *NbMgChl1* and *NbMgChl2* by all viruses delivering gRNAs, but the editing efficiency and mutations introduced varied among treatments (**Fig. 2, Table S3**). Among the TRV:tRNA-treated plants, the green sectors, present only on the first systemic leaves, were either wild type (*n* = 1 of 3 plants) or mosaic with 1 bp indels in the first and third target sites in both genes (*n* = 2 of 3 plants; **Fig. 2D**). The white sectors on the first and seventh systemic leaves of TRV:tRNA-treated plants harbored biallelic mutations: the first systemic leaves had 1 bp indels (*n* = 2 out of 3 plants) or 95 bp deletions in both loci (*n* = 1 plant); the seventh systemic leaves displayed biallelic 1 bp indels or 95 bp deletions in both homeologs (*n* = 2 out of 3 plants), or were mosaic at one locus and monoallelic for a 95 bp deletion at the second locus (*n* = 1 out of 3 plants). Larger deletions were more consistently observed among the TRV:mAtFT-treated plants (**Fig. 2E**). The green sectors from the first systemic leaves from TRV:mAtFT-treated plants were wild type (*n* = 2 out of 3 plants) or mosaic (*n* = 1 out 3 plants). Green sectors from the seventh leaves were present in only two TRV:mAtFT-treated plants: these tissues were wild type in one plant while the second plant harbored monoallelic deletions of 95 bp and 1 bp in both genes. The white sectors from the TRV:mAtFT-treated plants harbored biallelic mutations in both *NbMgChl* genes, with most samples having 95 or 113 bp deletions in each homeolog. Consistent with the observed phenotypes (**Fig. 1F**), the TRV:pri-miR399-treated plants showed spotty editing (**Fig. 2F**). The first (*n* = 3 out of 3 plants) and seventh (*n* = 3 out of 3 plants) systemic green leaves of TRV:pri-miR399-treated plants were wild type; the white sectors from the first systemic leaves were wild type or mosaic with a 1 bp indel; the white sectors from the seventh true leaves were wild type (*n* = 2 of 3 plants) or the homeologs harbored monoallelic 1 bp deletions in the first target sequence (*n* = 1 out of 3 plants). Mutations to the *pri-miR399* sequence altered the editing efficiency. The green tissues from TRV:mir-pro plants were generally wild type, but the white sectors from the first systemic leaves contained biallelic deletions of 113 or 95 bp or were mosaic for 1 bp deletions, and the white sectors from the seventh systemic leaves displayed biallelic (*n* = 2 out of 3 plants) or monoallelic (*n* = 1 out of 3 plants) 1 bp deletions in the first target sequence of both genes (**Fig. 2G**). The green leaves of TRV:mir-mimic-treated plants were wild type or mosaic (**Fig. 2H**). Only one TRV:mir-mimic-treated plant had photobleached tissues in the first systemic leaf and this harbored biallelic mutations of 95 bp in one gene and mosaic deletions in the *NbMgChl* homeolog. Both loci harbored biallelic (*n* = 2 of 3 plants) or monoallelic (*n* = 1 of 3 plants) 1 bp deletions in the first target sequence in the white sectors from the seventh systemic leaves (**Fig. 2H**). As expected, the uninoculated, TRV-treated, and TRV:NbMgChl-silenced plants showed no signs of editing of either homeolog (**Fig. 2B, C; Table S3**). These analyses showed that meristem editing by the viral vectors was influenced by the flanking mobility factors, with the *mAtFT* sequence generally contributing to larger deletions in both green and white systemic tissues compared to other treatments.

To test if virus delivery of gRNAs activated VIGS or if the incorporated mobility factors affected systemic virus movement, we quantified target gene expression and virus accumulation. The relative expression of *MgChl* and TRV transcripts in the white and green sectors from the systemic leaves of virus-treated and control plants was determined by RT-qPCR. As expected, the uniformly photobleached leaves from VIGS plants accumulated significantly less *NbMgChl* transcripts compared to uninoculated and TRV-treated controls (*p* < 0.05, *n* = 3; **Fig. 3A**). However, the *NbMgChl* transcripts were not significantly reduced in the white leaf sectors from other treatments when compared to the uninoculated and TRV controls (**Fig. 3A**). The green leaf sectors from TRV:mAtFT-treated plants accumulated significantly more *NbMgChl* transcripts than the uninoculated and TRV-treated controls, which may reflect compensation in highly photobleached plants. The accumulation of TRV RNA 2 did not differ significantly among virus treatments (**Fig. 3B**). These results suggested that all viruses replicated and moved systemically throughout the plants, and all viruses delivering gRNAs edited the intended targets.

**Fig. 3:**
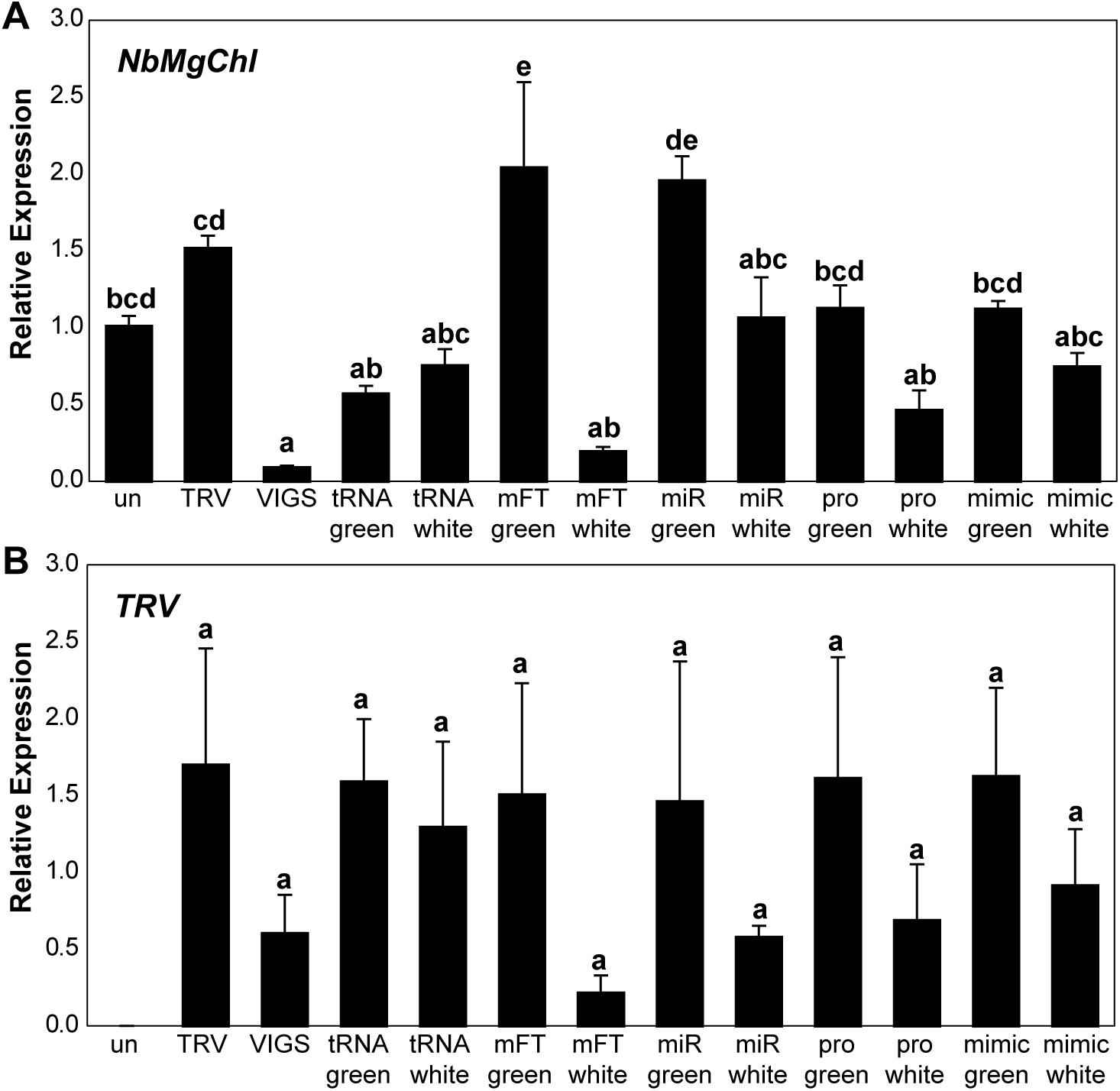
M**e**ristem **editing in *N. benthamiana* did not adversely affect the accumulation of targeted transcripts. (A)** The expression of the *NbMgChl* homeologs and **(B)** TRV is shown relative to *NbPP2a* and *NbEF1α* in the systemic green and white tissue sectors from uninoculated and virus-treated plants (*n* = 3 plants per treatment). The relative expressions of *NbMgChl* and *TRV* were compared to uninoculated and TRV controls, respectively. Error bars are means ± SE. The effects of treatment on gene expression were determined by ANOVA with post-hoc test by Tukey HSD, with significant differences at *p* < 0.05 indicated by lower case letters.

### Mobility factors differentially influence stable transmission of edited loci

To determine whether edits to the *NbMgChl* homeologs were successfully transmitted to the next generation, we examined the progeny from each treatment. While plants presenting extensive photobleaching produced fewer flowers and fruits, at least three fruits were harvested from each plant. The M1 varied extensively within and among treatments (**Fig. 4**). Some of the seedpods from the TRV:mAtFT-treated plants produced entirely white seeds, which failed to germinate (**Fig. S4**). The viable M1 from the seedpods collected from the TRV:tRNA- and TRV:mAtFT-treated plants ranged in color from green, pale green, to white (**Fig. 4D, E**), or were exclusively white (**Fig. 4F**). In contrast, the progeny from the remaining treatments, including the uninoculated, TRV-treated, and TRV:NbMgChl-silenced plants, were all green in color (**Fig. 4A-C, G-I**). These phenotypes suggested that, depending on the mobility factor used, edits were heritable but not necessarily clonal.

**Fig. 4:**
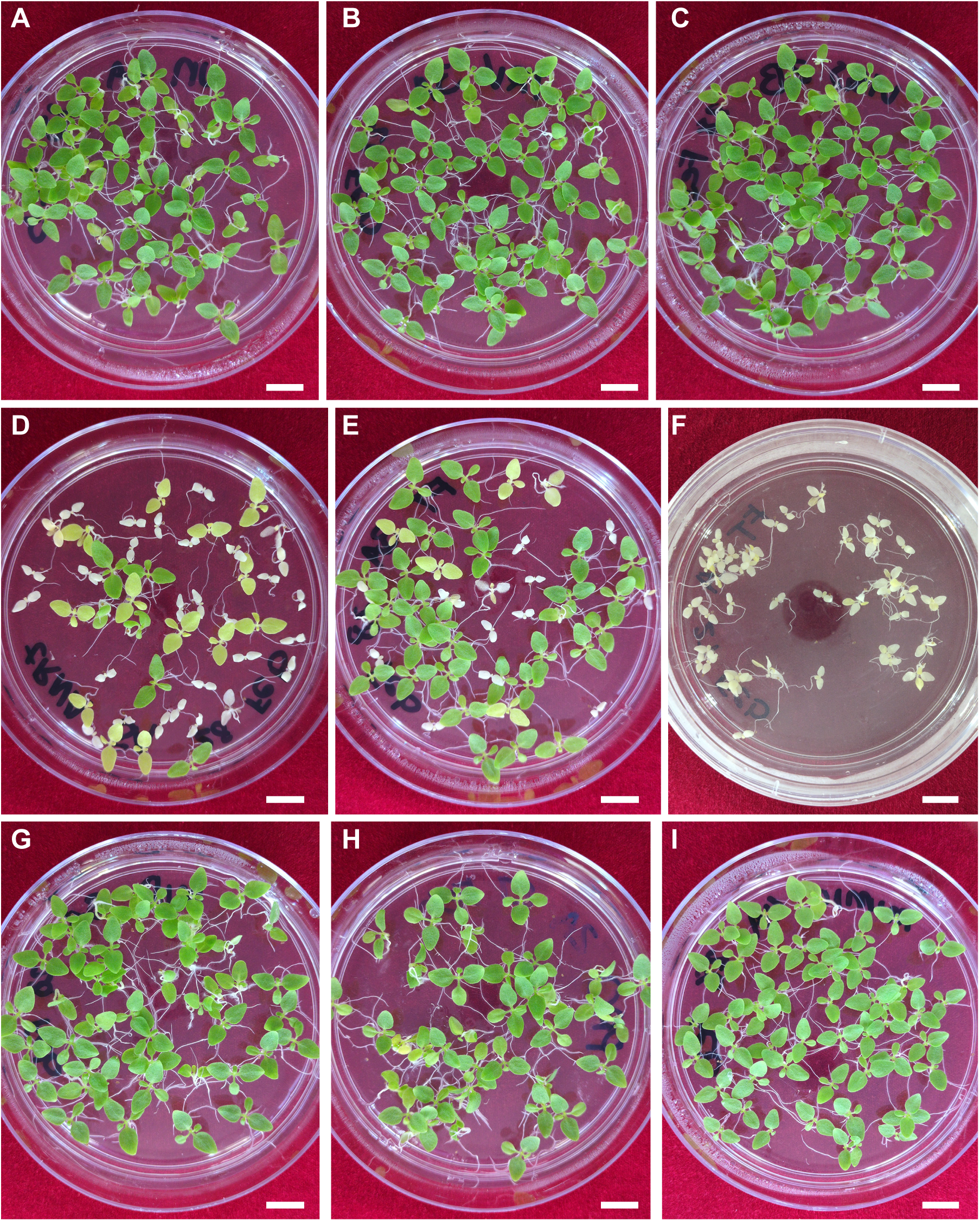
P**h**enotypic **evidence of heritable editing.** The progeny from individual seedpods from **(A)** uninoculated, **(B)** TRV-, **(C)** TRV:NbMgChl-, **(G)** TRV:miR399-, **(H)** TRV:mir-pro-, and **(I)** TRV:mir-mimic-treated plants were green. The progeny from **(D)** TRV:tRNA-, and **(E, F)** TRV:mAtFT-treated *N. benthamiana* showed segregating phenotypes. Some seedpods from TRV:mAtFT-treated plants produced only white colored seedings **(F)**. Scale bars are 1 cm, and petri dishes are the same size.

To further characterize these differences, we analyzed the genotypes of the M1 (**Fig. 5, Table S4**). Seedlings of each color were analyzed from individual fruits and from replicate plants.

**Fig. 5:**
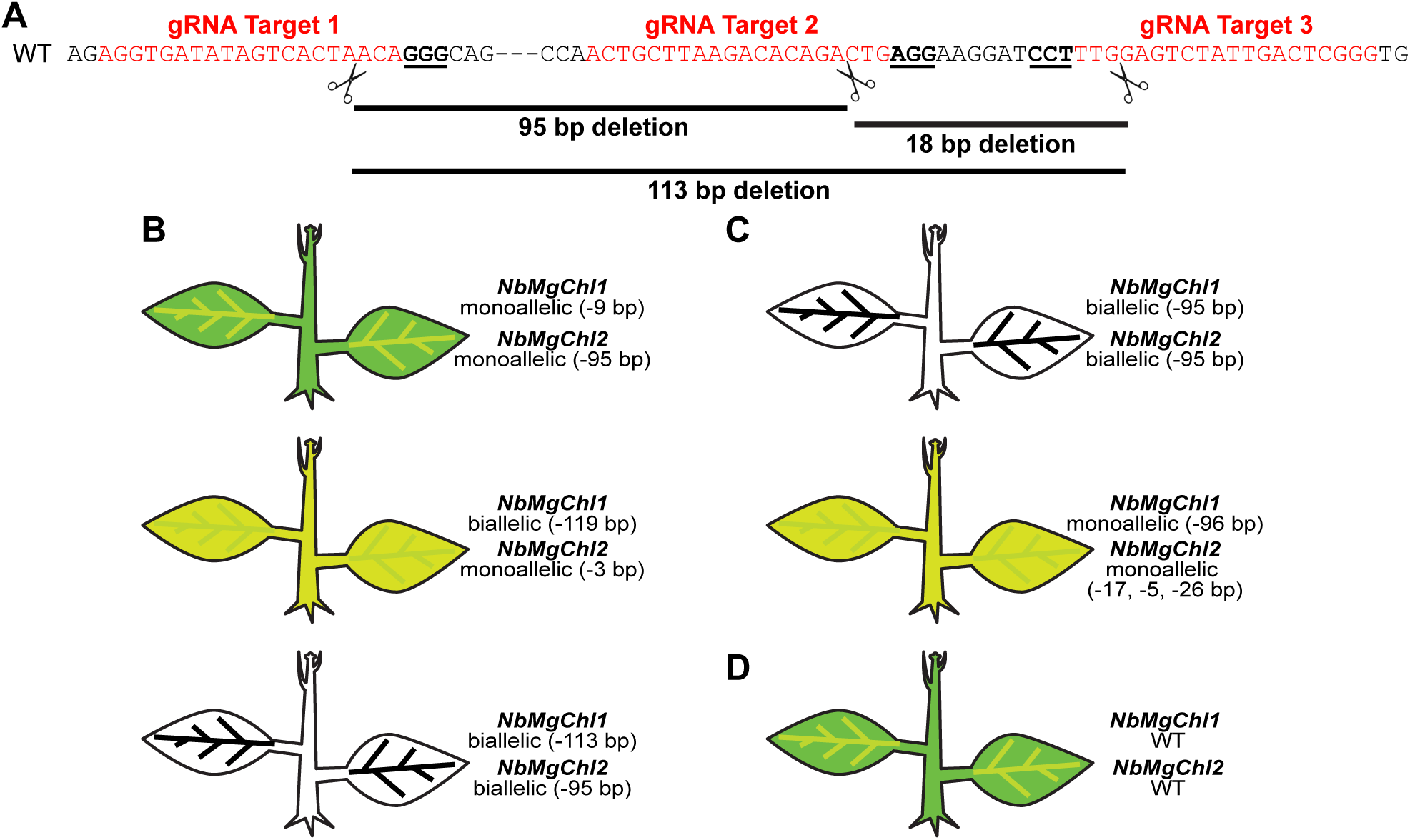
M**o**lecular **evidence of heritable editing. (A)** Shown is the region of the *NbMgChl1* and *NbMgChl2* genes targeted for editing. The gRNA targets are in red, with PAM sequences underlined. Predicted cut sites are represented with scissors, and the expected deletions resulting from multiple cuts are indicated. **(B)** The segregating progeny from TRV:tRNA-treated plants showed diverse edits (see also **Table S3**), with the number of wild-type alleles affecting seedling color. **(C)** The white M1 from TRV:mAtFT-treated plants harbored large biallelic mutations in both genes. **(D)** The green progeny from the uninoculated, TRV-, TRV:NbMgChl-, TRV:miR399-, TRV:mir-pro-, and TRV:mir-mimic-treated plants were wild type.

Various mutations were identified in the green seedlings from the TRV:tRNA-treated plants. Most frequently, the green seedlings harbored monoallelic mutations in both genes (*n* = 3 out of 8 green seedlings). Green seedlings also demonstrated monoallelic mutations in one gene with the second gene being mosaic or wild type (*n* = 2 out of 8 green seedlings); biallelic mutations in one gene with the second gene wild type (*n* = 1 out of 8 green seedlings); biallelic mutations in both genes (*n* = 1 out of 8 green seedlings); or were wild type (*n* = 1 of 8 green seedlings). The pale green seedlings from the TRV:tRNA-treated plants exhibited more consistent genotypes: plants harbored biallelic mutations in one gene and monoallelic mutations in the second gene (*n* = 7 out of 8 pale green seedlings); and one pale green seedling harbored monoallelic 95 bp deletions in each gene (*n* = 1 out of 8 pale green seedlings). The white seedlings from this treatment typically had large, biallelic deletions (- 95 or -113 bp) in both genes (*n* = 6 out of 8 white seedlings) or biallelic mutations in one gene with the wild-type copy of the second gene (*n* = 2 out of 8 white seedlings). These results suggested that the homeologous *MgChl* genes were partially redundant, and the dosage of wild-type gene copies affected seedling phenotypes. The white seedlings from the TRV:mAtFT-treated plants harbored large and biallelic deletions (-95 bp) in both *NbMgChl* genes (*n* = 8 out of 8 white seedlings). The one pale green seedling analyzed from TRV:mAtFT-treated plants harbored a monoallelic 96 bp deletion in one gene while the second *NbMgChl* gene had monoallelic mutations in all three target sequences. All remaining treatments produced green seedlings, which were genotypically wild type. Our findings demonstrated that not all of the viruses edited the germline. However, complex and heritable editing was possible when *tRNA^Gly^* and *mAtFT* sequences were incorporated in the TRV vectors delivered to *N. benthamiana*.

### Extending TRV delivery of editing tools to an emerging oilseed crop

Despite its limited amenability to standard transformation procedures, there are no reports to date of virus-based transient genetic manipulation in pennycress. We hypothesized that our virus tools could be adapted to edit pennycress meristems and advance functional genomics. To test this, we sought to first determine whether TRV was infectious in pennycress using VIGS assays.

The pennycress ortholog (Thlar.0013s1122, *TaMgChl*) of the Arabidopsis *MgChl* (At5G45930) was identified through BLASTp searches of the *Thlaspi arvense* proteome. A 300-bp fragment of *TaMgChl* was used to construct the silencing vector TRV:TaMgChl. The silencing vector and the TRV control were delivered to wild-type pennycress via *Agro*-infiltration into leaves or vacuum-infiltration into the whole seedlings; uninoculated controls remained uninfiltrated. The pennycress plants treated with TRV:TaMgChl, using either delivery method, presented the expected photobleached phenotypes (**Table S5**). Unlike the highly consistent phenotypes observed among the *N. benthamiana* plants treated with TRV:NbMgChl, photobleaching in the treated pennycress plants was variable. The onset of photobleaching varied, with phenotypes visible as early as 13 dpi or as late as 25 dpi. Photobleaching typically extended incompletely from the mid-rib to the leaf periphery (**Fig. 6C**). The duration of silencing was inconsistent among replicate plants: white silicles were occasionally observed (**Fig. 6D**), and seemingly unsilenced plants unexpectedly produced racemes with a few white internodes before newer green growth resumed (**Fig. 6E**). The TRV-treated and uninoculated pennycress plants remained green and healthy (**Fig. 6A, B**). Though inconsistent, these experiments demonstrated that TRV was infectious in pennycress, and delivery of TRV via *Agro*-infiltration of leaves and vacuum infiltration of whole seedlings achieved VIGS (**Table S5**).

**Fig. 6:**
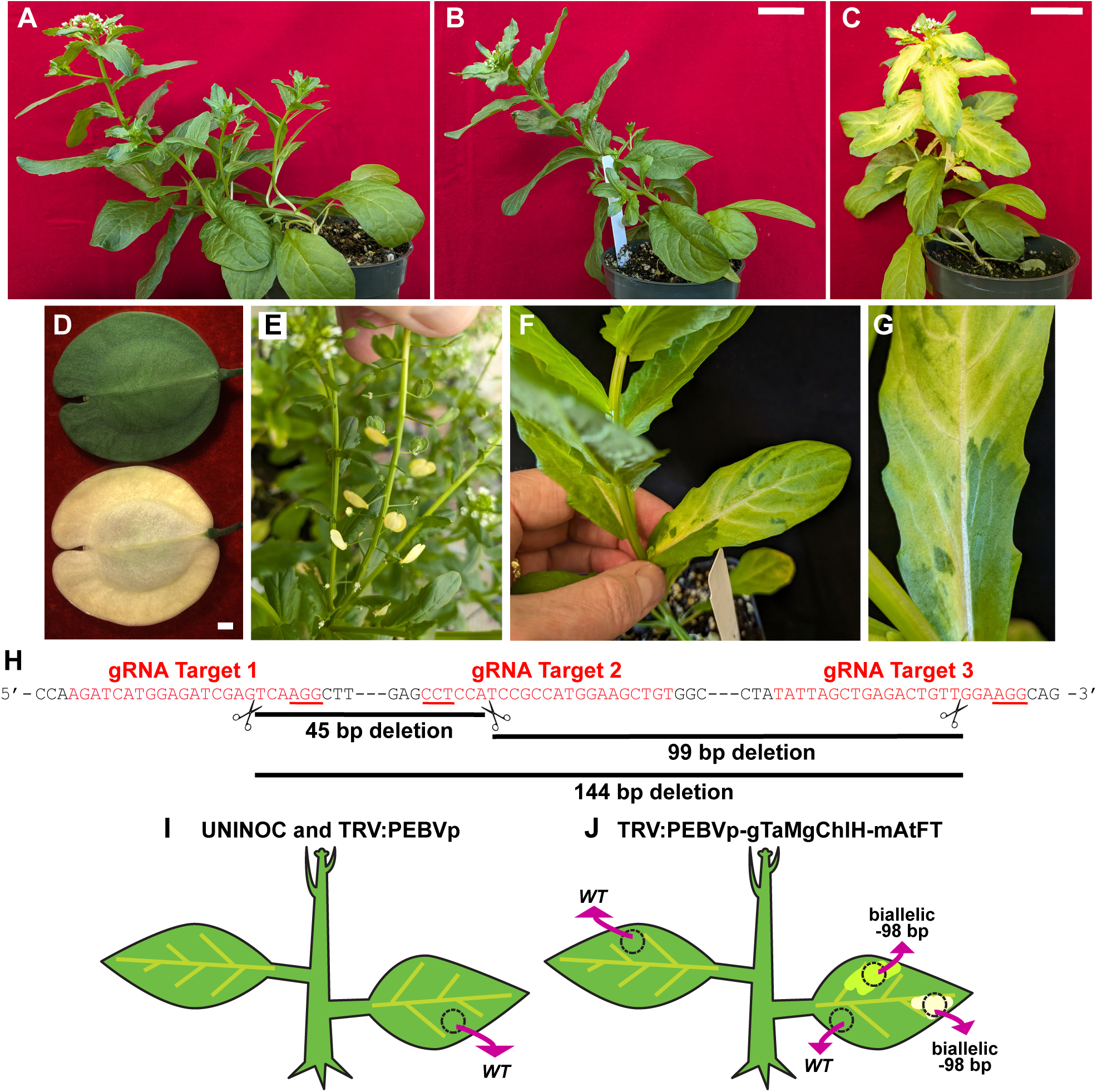
E**x**tending **virus-based tools for transient gene manipulation and meristem editing in pennycress. (A)** Uninoculated and **(B)** TRV-treated pennycress plants. **(C-E)** TRV:TaMgChl-treated pennycress shows systemic silencing in leaves and stems **(C)**, silicles **(D)** and inconsistently in inflorescences **(E)**. Pots in **(A-C)** were the same size, and scale bars in **(B, C)** are 5 cm and in **(D)** is 1 mm. (F, G) TRV:mAtFT-treated pennycress produced sectored systemic leaves. **(H)** The sequence of the TaMgChl gene targeted for editing. The gRNA targets are shown in red, with PAM sites underlined. The predicted cut sites are represented with scissors, and the expected deletions resulting from multiple cleavages are indicated. **(I)** Leaves from uninoculated and TRV-treated pennycress plants were wild type. **(J)** The pale-green and white sectors present in a systemic leaf from a TRV:mAtFT-treated plant harbored targeted biallelic mutations.

To test if TRV could be adapted to edit pennycress meristems, we constructed a new virus targeting *TaMgChl* for editing. Using CRISPR RGen Tools, we identified three gRNAs to target *TaMgChl* with specificity. Based on the results from *N. benthamiana*, the *mAtFT* mobility factor was selected as the most promising element to achieve meristem editing. The gRNAs were introduced from overlapping oligonucleotides for seamless assembly with the *mAtFT* mobility factor in TRV:PEBVp, as per the design strategies used in *N. benthamiana* (**Fig. S1A, C**). The editing construct, TRV:PEBVp:gTaMgChl-mAtFT, the TRV:TaMgChl silencing construct, and TRV:PEBVp were delivered to *35S:Cas9* pennycress plants using leaf- and vacuum-infiltration methods; the uninoculated controls remained uninfiltrated. Across multiple independent experiments, a total of three out of 54 TRV:PEBVp:gTaMgChl-mAtFT-treated plants each produced a single, sectored leaf (**Fig. 6F, G, Table S5**). The sectored leaves appeared similar to the silencing observed in the TRV:TaMgChl-treated pennycress but sectoring did not continue to new tissues; these phenotypes were not observed in the control treatments. To determine if the sectors reflected targeted editing events, genomic DNA was isolated from the dark green, pale green, and white sectors, as well as from the next, non-sectored systemic leaf, and prepared for amplicon sequencing. As controls, genomic DNA was collected from the TRV:PEBVp-treated and uninoculated plants. Amplicon sequencing identified targeted editing of *TaMgChl*: the pale green and white sectors from the leaves of TRV:PEBVp:gTaMgChl-mAtFT-treated plants harbored monoallelic or biallelic 98 bp deletions (**Fig. 6H, J, Table S6**). The dark green sectors from the same leaves were wild type (**Fig. 6H, J; Table S6**). The leaf immediately following the sectored leaf was mosaic with three bp deletions (**Table S6**). The green leaves from the uninoculated and TRV:PEBVp-treated pennycress plants were wild type (**Fig. 6I, Table S6**). Our findings demonstrated successful meristem editing of a non-model plant using TRV.

## DISCUSSION

Meristem-based gene editing with plant viruses is an exciting approach to overcome the transformation barriers that limit functional analyses in many plants. To date, there are reports of a few cloned viruses that achieve somatic and heritable edits in *Cas9*-expressing plants (Beernink et al., 2022; Ellison et al., 2020; Kang et al., 2025; Li et al., 2021; Mei et al., 2019; Nagalakshmi et al., 2022b; Uranga et al., 2023; Uranga et al., 2021). More recently, TRV was shown to deliver the smaller endonuclease ISYmu1 with a single RNA guide to Arabidopsis, potentially extending meristem editing to any genetic background (Weiss et al., 2025). The roles of mobility factors in delivering editing tools to plant meristems from a mobile virus are not straightforward. While incorporating *mAtFT* dramatically improved heritable gene editing in *N. benthamiana* (Ellison et al., 2020), this success did not extend to editing *Cas9-*expressing Arabidopsis (Nagalakshmi et al., 2022a). Incorporating tRNAs, *mAtFT*, or the non-coding sequence from the wheat (*Triticum aestivum*) *FT* ortholog, *mTaFT*, in Barley stripe mosaic virus yielded heritable edits in *Cas9*-transgenic wheat (Li et al., 2021), but not in *Cas9*-transgenic *N. benthamiana*. (Beernink et al., 2022; Uranga et al., 2021). Moreover, the addition of the same mobility factors into Barley stripe mosaic virus, Foxtail mosaic virus, and Potato virus X impaired or promoted germline edits in the species tested (Beernink et al., 2022; Li et al., 2021; Mei et al., 2019; Uranga et al., 2021)(Kang et al., 2025). This variability underscored the need to identify a range of mobile RNA sequences for broad utility.

Here, we constructed a series of TRV vectors for multiplexed gene editing. The modular nature of our vectors facilitated the easy exchange of gRNA targets and/or mobility factors, providing a foundation to optimize virus design. We directly compared the impacts of different candidate mobility factors on editing visible marker genes. When delivered to *Cas9*-expressing *N. benthamiana*, all of the TRV viruses harboring gRNAs resulted in edits in the target genes in somatic tissues. Among the treatments tested, the addition of the *tRNA^Gly^* and *mAtFT* sequences profoundly affected somatic editing of the *NbMgChl1* and *NbMgChl2* genes, resulting in large targeted biallelic deletions (**Fig. 2**). The presence of either of these mobility factors enhanced the transmission of edited alleles to the next generation, with the M1 collected from TRV:mAtFT-treated plants prominently displaying strong loss-of function phenotypes and biallelic mutations in both target genes. Unexpectedly, these experiments demonstrated that the *NbMgChl1* and *NbMgChl2* gene products function in a dose-dependent manner: at least one copy of each gene is needed for green pigmentation (**Fig. 5**). We hypothesized that additional RNA secondary structures could extend the range of meristem editing from TRV. When tested in *N. benthamiana*, the *pri-miR399* and variants resulted in targeted somatic edits, but edits were not conveyed to the next generation. We recognized that the secondary-structure rich *pri-miR399* sequence could have been processed early in viral replication, undermining delivery of gRNAs. However, the variants, which were designed to retain the RNA secondary structure, performed similarly. These findings suggest that secondary structure alone is insufficient for gRNA delivery. It would be intriguing to compare when and where edits occurred from these different viruses. Robust somatic and heritable editing, as observed with TRV:tRNA and TRV:mAtFT, may result from early edits to stem cells. In contrast, editing events occurring later in the emergence of the leaf primordium would be expected to affect only some somatic tissues, as observed in plants treated with TRV:pri-miR399, TRV:mir-pro, and TRV:mir-mimic.

*N. benthamiana* is a valued tool for plant-virus interactions. With a deletion in *RNA DEPENDENT RNA POLYMERASE 1* (Bally et al., 2018; Yang et al., 2004) and genomic losses of additional components of the host silencing machinery (Wang et al., 2024), *N. benthamiana* is hyper-susceptible to virus infections. While these attributes enabled the quantitative dissection of component elements tested in our TRV vectors, it was imperative to extend findings from this ideal host to non-model systems. Here, we presented the first demonstration of virus-based genetic manipulation in pennycress. While the VIGS assays achieved the expected photobleached phenotype, the inconsistency observed over multiple independent experiments suggested that TRV may not be the ideal virus to advance pennycress functional genomics. Notwithstanding, we leveraged the strong *mAtFT* mobility factor to deliver editing tools from TRV to pennycress meristems, and produced large, targeted biallelic mutations in systemic leaves. Targeted edits were limited and we did not observe phenotypes in silicles that were consistent with intended edits, and these outcomes likely reflect the suboptimal performance of TRV in pennycress. Collectively, our findings highlight the promise of virus-based meristem editing for accelerating trait development in non-model crops and underscore the need for improved viral platforms to achieve consistent and heritable outcomes.

## SUPPORTING INFORMATION

**Table S1:** Primer sequences.

**Table S2:** Sequences of mobility factors incorporated in TRV RNA 2.

**Table S3:** Amplicon sequencing identifies somatic edits in *N. benthamiana*.

**Table S4:** Amplicon sequencing identifies heritable edits in *N. benthamiana*.

**Table S5:** Summary of VIGS and meristem editing experiments in pennycress.

**Table S6:** Amplicon sequencing identifies somatic edits in pennycress.

**Fig. S1:** Construction of viruses for gene editing.

**Fig. S2:** Silencing *NbMgChl1* and *NbMgChl2* by VIGS yields the expected photobleached phenotype.

**Fig. S3:** RNAfold predicted RNA secondary structures.

**Fig. S4:** Meristem editing impacted seed color and viability in treated *N. benthamiana* plants.

## DATA AVAILABILITY

The author responsible for the distribution of data and materials integral to the findings presented in this article and in accordance with this journal’s policies on data sharing is RCM.

## AUTHOR CONTRIBUTIONS

RCM designed experiments; MA designed rational mutations in *pri-miR399* and generated the *Cas9*-expressing pennycress line used; KTC, HH, and RCM conducted experiments in *N. benthamiana*; YTL and RCM conducted experiments in pennycress; RCM analyzed the data and wrote the manuscript; all authors commented on the manuscript.

## ACKNOWLEDGEMENTS

We are grateful to Dr. Dan Voytas (University of Minnesota) for generously providing *Cas9*-expressing *N. benthamiana*, and to Dr. Savio S. Ferreira (University of North Texas) for generously sharing the *Cas9*-expressing pennycress line. We thank Dr. Guadalupe Lopez Puc (CIATEJ) for her assistance with early VIGS experiments in *N. benthamiana*.

