## Supporting Information - Table and Figure Legends for "Mobility-enhanced virus vectors enable meristem genome editing in model and crop plants"

**Table S1: Primer sequences.** Oligonucleotides used for cloning gRNAs in viral vectors, amplicon sequencing, and RT-qPCR are shown in separate tabs. Primers for RT-qPCR do not distinguish between homeologous genes.

**Table S2: Sequences of mobility factors incorporated in TRV RNA 2.** The *tRNA$^{Gly}$* (Xie et al., 2015) and *mAtFT* (Ellison et al., 2020) sequences are shown. The primary microRNA sequence derived from the wild-type *pri-miR399b* sequence is shown. Modified nucleotides are indicated in lower-case red font. Mature *miR399b* and *miR399b** sequences are underlined.

**Table S3: Amplicon sequencing identifies somatic edits in *N. benthamiana*.** Molecular analyses of editing from each leaf sample are shown (sample indicated in column A). As described in(Xue and Tsai, 2015), each target, with forward and reverse reads, is indicated in column C, with the target sequence included in column D. The main edited sequence for each target is in column E (dashes in the sequence represent deletions compared to the target sequence). The total hits (column F) indicate the total number of reads matching the *NbMgChl1* and *NbMgChl2* target sequences in the file. The sub-hits in column G shows the number of reads matching each target, with the proportion of wild-type and edited hits included in columns H and I. The pattern (column J) indicates the nature and size of the edit. Samples are color-coded for easier viewing.

**Table S4: Amplicon sequencing identifies heritable edits in *N. benthamiana*.** The amplicon sequencing results of the progeny plants are shown. The sample name (column A) represents the parent treatment (eg. uninoc 3A), color of the $M_1$ seedling (eg. green), and independent progeny analyzed (1 - 4). The sequencing results are organized as in **Table S3**.

**Table S5: Summary of VIGS and meristem editing experiments in pennycress.** Details of five independent VIGS and editing experiments in pennycress. The method of inoculation and age of inoculated plants are shown in column A, with treatment (column B), number of plants treated (column C), and the number of plants showing the indicated phenotype (column D), included. Experiments are color-coded for easier viewing.

**Table S6: Amplicon sequencing identifies somatic edits in pennycress.** Molecular analyses of editing are shown. The sample name (column A) indicates the treatment, replicate plant, and tissue sector analyzed. The remaining columns are as in **Table S3** and **Table S4**.

**Fig. S1: Construction of viruses for gene editing. (A)** TRV is a bipartite RNA virus. The TRV RNA 1 genome encodes the RNA-dependent RNA polymerase (RdRP), the movement protein (MP), and 16 K non-structural protein. TRV RNA 2 encodes the coat protein (CP). The Pea early browning virus coat protein promoter (PEBVp) was introduced upstream of the multiple cloning site, as described (McGarry and Ayre, submitted). **(B)** The TRV:tRNA virus construction is illustrated. The three gene editing cassettes, each comprised of the gRNA (red box), scaffold (blue box), and $tRNA^{Gly}$ (green box), were assembled. The gRNAs were introduced on overlapping oligonucleotides, and each cassette was PCR amplified. The three PCR reactions were pooled, column-purified, and digested with restriction endonucleases as shown. These were assembled as ligated into TRV RNA 2 harboring the PEBVp ("modified TRV RNA 2") shown in **(A)**. **(C)** The TRV:mAtFT virus construction is shown. The assembly was as in **(B)**. **(D)** The TRV:miR399 virus was constructed from three separately synthesized plasmids as described in the **Materials and Methods**. The assembly of TRV:mir-pro and TRV:mir-mimic was as in **(D)**.

**Fig. S2: Silencing *NbMgChl1* and *NbMgChl2* by VIGS yields the expected photobleached phenotype. (A)** Uninoculated, **(B)** TRV-treated, and **(C)** TRV:NbMgChl-silenced plants. Plants are shown at 10 dpi. Scale bars are 5 cm.

**Fig. S3: RNAfold predicted RNA secondary structures of rationally modified *pri-miRNA399b*-based mobility factors compared to *mAtFT*. (A)** *Pri-miRNA399b* predicted structure. The boxed region indicates the mature *miRNA399b* sequence. Arrows show the locations of rational sequence modifications introduced to prevent the processing of the *pri-miRNA* into a mature sequence, yielding the *mir-pro* mobility factor. **(B)** The *mAtFT* predicted structure. **(C)** the *mir-mimic* predicted structure. The minimum free energy structure drawings encoding base-pair probabilities are color-coded according to the scale shown.

**Fig. S4: Meristem editing impacted seed color and viability in treated *N. benthamiana* plants. (A)** A white seedpod obtained from a *N. benthamiana* plant treated with TRV:mAtFT yielded brown seeds. **(B)** A sectored seedpod obtained from a replicate plant treated with TRV:mAtFT produced all white seeds. The white seeds failed to germinate. Scale bars are 500 μm.

Ellison, E.E., Nagalakshmi, U., Gamo, M.E., Huang, P.-j., Dinesh-Kumar, S. and Voytas, D.F. (2020) Multiplexed heritable gene editing using RNA viruses and mobile single guide RNAs. Nat. Plants 6, 620–624.

Xie, K., Minkenberg, B. and Yang, Y. (2015) Boosting CRISPR/Cas9 multiplex editing capability with the endogenous tRNA-processing system. Proc Natl Acad Sci USA 112, 3570–3575.

Xue, L.-J. and Tsai, C.-J. (2015) AGEseq: Analysis of genome editing by sequencing. Mol Plant 8, 1428–1430.
