## Supplementary material for "Mobility-enhanced virus vectors enable meristem genome editing in model and crop plants": Fig. S1

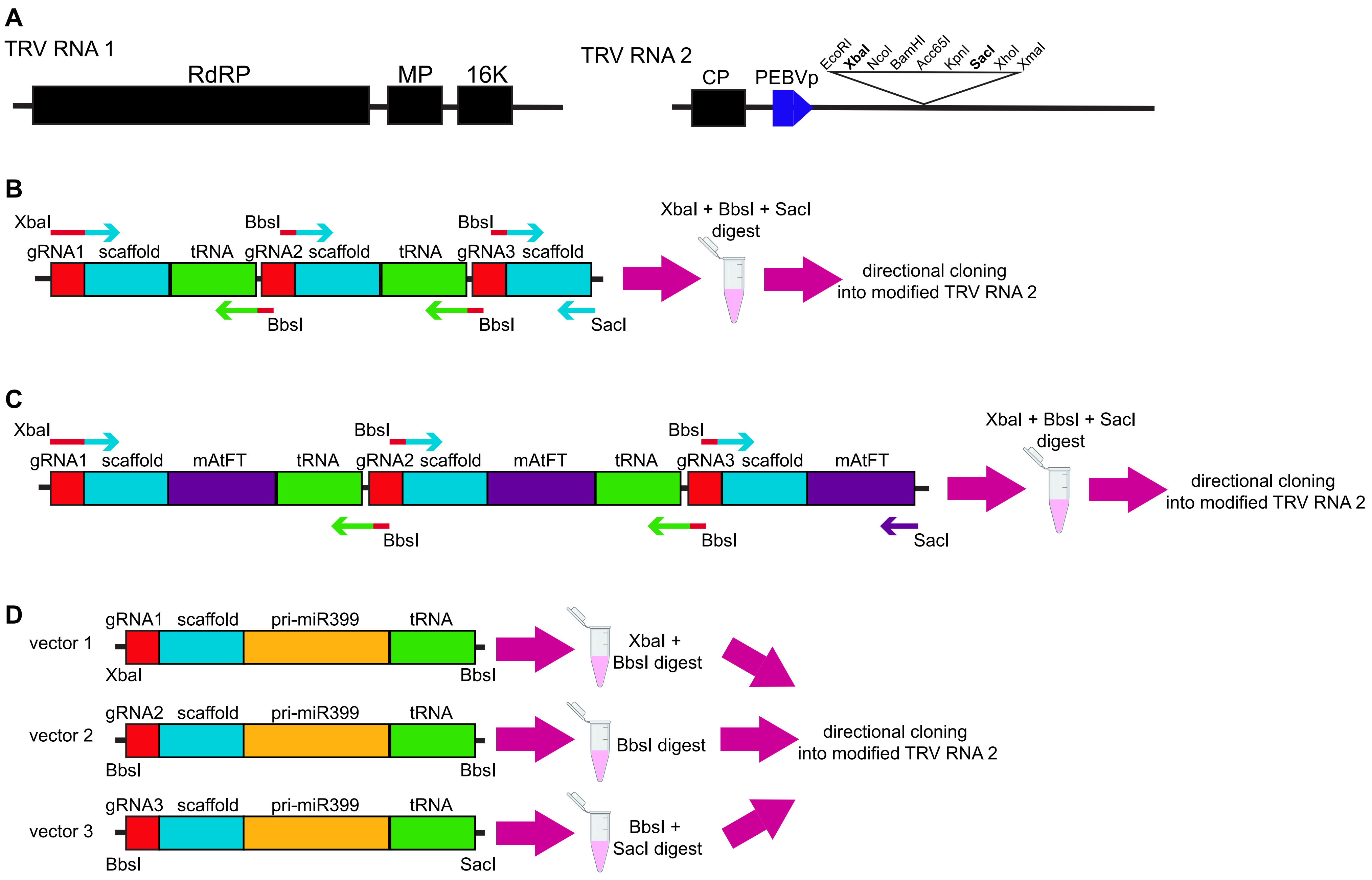


**Fig. S1: Construction of viruses for gene editing. (A)** TRV is a bipartite RNA virus. The TRV RNA 1 genome encodes the RNA-dependent RNA polymerase (RdRP), the movement protein (MP), and 16 K non-structural protein. TRV RNA 2 encodes the coat protein (CP). The Pea early browning virus coat protein promoter (PEBVp) was introduced upstream of the multiple cloning site, as described (McGarry and Ayre, submitted). **(B)** The TRV:tRNA virus construction is illustrated. The three gene editing cassettes, each comprised of the gRNA (red box), scaffold (blue box), and $tRNA^{Gly}$ (green box), were assembled. The gRNAs were introduced on overlapping oligonucleotides, and each cassette was PCR amplified. The three PCR reactions were pooled, column-purified, and digested with restriction endonucleases as shown. These were assembled as ligated into TRV RNA 2 harboring the PEBVp ("modified TRV RNA 2") shown in **(A)**. **(C)** The TRV:mAtFT virus construction is shown. The assembly was as in **(B)**. **(D)** The TRV:miR399 virus was constructed from three separately synthesized plasmids as described in the **Materials and Methods**. The assembly of TRV:mir-pro and TRV:mir-mimic was as in **(D)**.
