## Supplementary material for "Mobility-enhanced virus vectors enable meristem genome editing in model and crop plants": Fig. S2

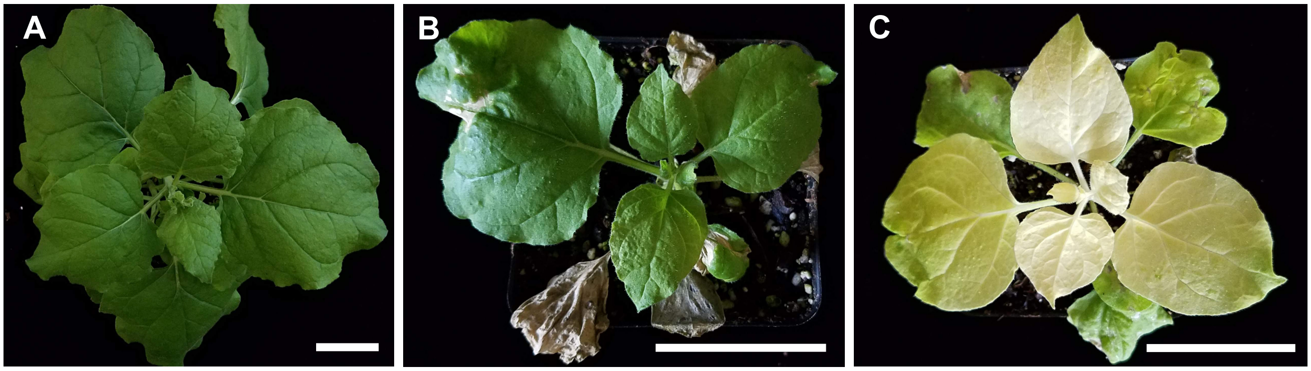


**Fig. S2: Silencing *NbMgChl1* and *NbMgChl2* by VIGS yields the expected photobleached phenotype. (A)** Uninoculated, **(B)** TRV-treated, and **(C)** TRV:NbMgChl-silenced plants. Plants are shown at 10 dpi. Scale bars are 5 cm.
