## Supplementary material for "Mobility-enhanced virus vectors enable meristem genome editing in model and crop plants": Fig. S3

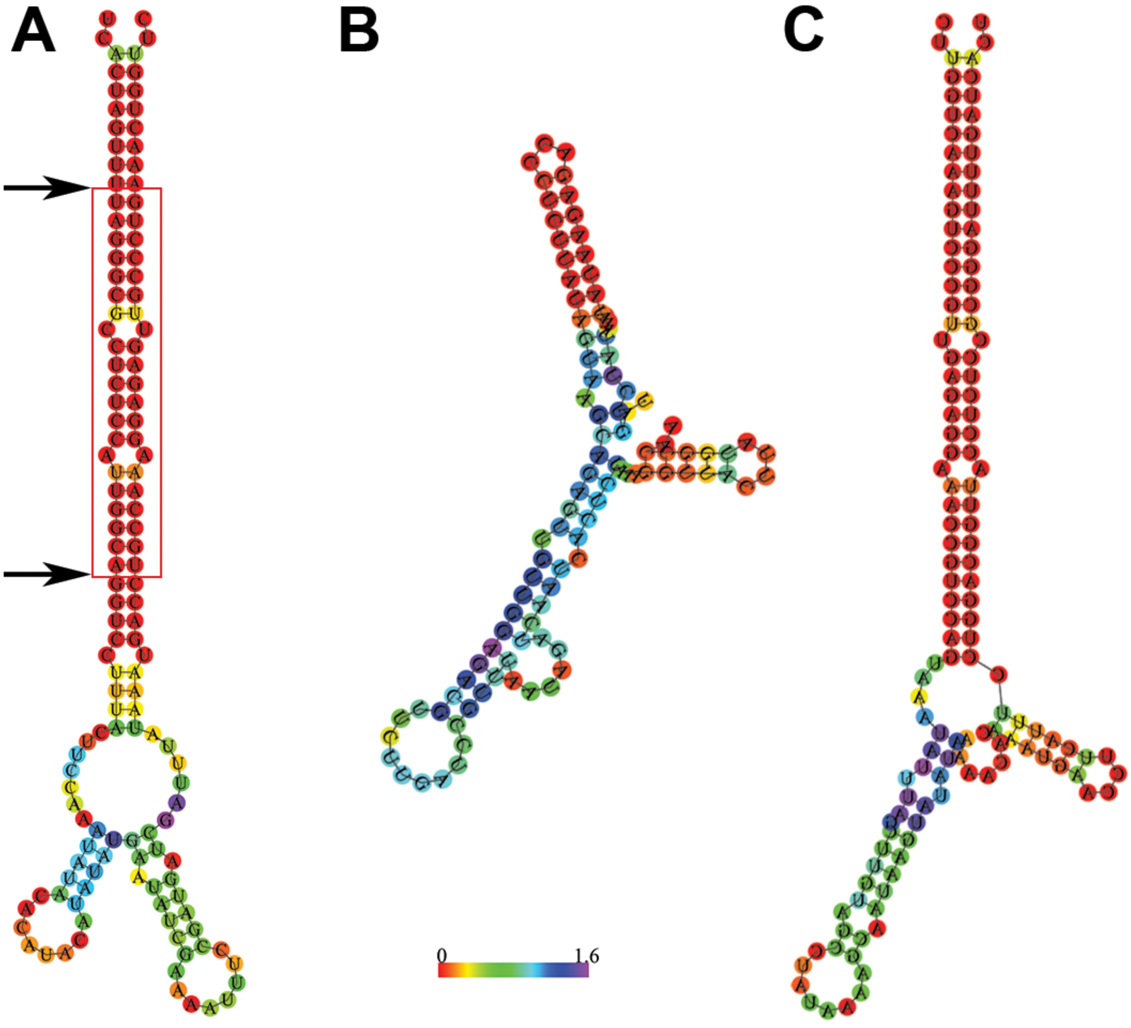


**Fig. S3: RNAfold predicted RNA secondary structures of rationally modified *pri-miRNA399b*-based mobility factors compared to *mAtFT*. (A)** *Pri-miRNA399b* predicted structure. The boxed region indicates the mature *miRNA399b* sequence. Arrows show the locations of rational sequence modifications introduced to prevent the processing of the *pri-miRNA* into a mature sequence, yielding the *mir-pro* mobility factor. **(B)** The *mAtFT* predicted structure. **(C)** the *mir-mimic* predicted structure. The minimum free energy structure drawings encoding base-pair probabilities are color-coded according to the scale shown.
