## Supplementary material for "Mobility-enhanced virus vectors enable meristem genome editing in model and crop plants": Fig. S4

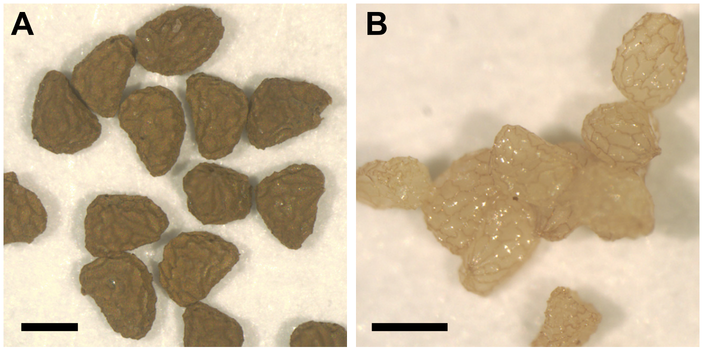


**Fig. S4: Meristem editing impacted seed color and viability in treated *N. benthamiana* plants.** **(A)** A white seedpod obtained from a *N. benthamiana* plant treated with TRV:mAtFT yielded brown seeds. **(B)** A sectored seedpod obtained from a replicate plant treated with TRV:mAtFT produced all white seeds. The white seeds failed to germinate. Scale bars are 500 µm.
