## Supplementary material for "Mobility-enhanced virus vectors enable meristem genome editing in model and crop plants": Table S2

**Table S2:** **Sequences of mobility factors incorporated in TRV RNA 2.**

| **Mobility Factor** | **RNA sequence (5ʹ)** |
| --- | --- |
| tRNA^Gly^ | AACAAAGCACCAGTGGTCTAGTGGTGGAATAGTACCCTGCCACGGTACAGACCCGGGTTCGATTCCCGGCTGGTGCA |
| mAtFT | TAGTCTATAAATATAAGAGACCCTCTTATAGTAAGCAGAGTTGTTGGAGACGTTCTTGATCCGTTTAATAGATCAATCACTCTAAAGGTTACTTATGGCCAA |
| Pri-miR399b (wt) | UCACUAGUUUUAGGGCGCCUCUCCAUUGGCAGGUCCUUUACUUCCAAAUAUACACAUACAUAUAUGAAUAUCGAAAAUUUCCGAUGAUCGAUUUAUAAAUGACCUGCCAAAGGAGAGUUGCCCUGAAACUGGUUC |
| Pri-miR399b-mimic variant (“mir-mimic”) | UCACUAGUUUUAGGGCGCCUCUCCAUUGGCAGGUCCUUUACUUCCAAgUAaACACAaAaAUAUAUGAAUAaCGAAAAUaUCCGAUGuUCGAUUUAUAAAUGACCUGCCAAAGGAGAGUUGCCCUGAAACUGGUUC |
| Pri-miR399b-processing variant (“mir-pro”) | UCACUAGUUUUAGGGCGCCUCUCCAUUGGCAGGUCaUUUAuaaauugAUcauuggAaACAUAUAUGAAUAUCGAAAAUUUCCGAUGAUCGAUUUAUAAAUGACCUGCCAAAGGAGAGUUGCCCUGAAACUGGUUC |

The *tRNA^Gly^* (Xie et al., 2015) and *mAtFT* (Ellison et al., 2020) sequences are shown. The primary microRNA sequence derived from the wild-type *pri-miR399b* sequence is shown. Modified nucleotides are indicated in lower-case red font. Mature *miR399b* and *miR399b** sequences are underlined.
